# Inhibitory hippocampo-septal projection controls locomotion and exploratory behavior

**DOI:** 10.1101/2021.09.17.460838

**Authors:** Yuh-Tarng Chen, Jun Guo, Wei Xu

## Abstract

Although the hippocampus is generally considered a cognitive center for spatial representation, learning and memory, increasing evidence supports its roles in regulation of locomotion. However, the neuronal mechanisms of hippocampal regulation of locomotion and exploratory behavior remain unclear. Here we found that the inhibitory hippocampo-septal projection bi-directionally controls locomotion speed of mice. Pharmacogenetic activation of these septum-projecting interneurons decreased locomotion and exploratory behavior. Similarly, activation of the hippocampus-originated inhibitory terminal in the medial septum reduced locomotion. On the other hand, inhibition of the hippocampus-originated inhibitory terminal increased locomotion. The locomotion-regulative roles were specific to the septal projecting interneurons as activation of hippocampal interneurons projecting to the retrosplenial cortex did not change animal locomotion. Therefore, this study reveals a specific long-range inhibitory output from the hippocampus in the regulation of animal locomotion.

## Introduction

Freely moving and exploring in an environment are essential skills for construction of a neuronal representation of the world and survival. Locomotion is regulated by brain motor systems consisting of the brain regions such as the motor cortex, basal ganglions, thalamus, spinal cord and so on (Ferreira-Pinto, Ruder, Capelli, & Arber, 2018). Normally the hippocampus is not considered as a component of this system, but rather as a cognitive center that integrates multi-modal sensory information and their spatial/temporal relations to construct a representation and memory of the world (Eichenbaum, 2004; Buzsaki & Moser, 2013). However, accumulating evidence indicates a direct role of the hippocampus in regulation of locomotion.

Firstly, hippocampal neuronal activities, especially that in the septal hippocampus, are associated with locomotion. Most noticeably, the activity level of some hippocampal neurons directly reflect the speed of animals (McNaughton, Barnes, & O’Keefe, 1983; Geisler, Robbe, Zugaro, Sirota, & Buzsaki, 2007; Gois & Tort, 2018; Iwase, Kitanishi, & Mizuseki, 2020). Interestingly, many of these “speed cells” were found to be inhibitory GABAergic “interneurons” (Gois & Tort, 2018; Iwase et al., 2020). Secondly, lesions or pathogenic damages of the hippocampus are frequently accompanied by alterations in locomotion (Sams-Dodd, Lipska, & Weinberger, 1997; Katsuta, Umemura, Ueyama, & Matsuoka, 2003; Godsil, Stefanacci, & Fanselow, 2005; White, Whitaker, & White, 2006). Thirdly, direct functional manipulations of the hippocampal neuronal activities with optogenetic or pharmacogenetic stimuli altered animal locomotion (Bender et al., 2015; Wolff et al., 2018). However, the studies of hippocampal roles in locomotion were full of contradictory results. For example, electrolytic lesion of the hippocampus caused hyperlocomotion (Douglas & Isaacson, 1964) and enhanced the locomotion-stimulating effect of amphetamine (Swerdlow, Halim, Hanlon, Platten, & Auerbach, 2001); while aspiration lesion of the hippocampus did not produce similar impacts (Douglas & Isaacson, 1964). Furthermore, unlike in the work mentioned above, some pharmacological, optogenetics, or pharmacogenetic treatments of the hippocampus failed to change animal locomotion activity (Zhang, Bast, & Feldon, 2002; Degoulet, Rouillon, Rostain, David, & Abraini, 2008; Goshen et al., 2011; Lopez et al., 2016; Bian et al., 2019)

The contradictions may arise from the anatomical and functional heterogeneity of the hippocampus. The hippocampus has multiple functional divisions along its longitudinal and transversal axis (Soltesz & Losonczy, 2018). It consists of numerous neuronal types including both excitatory glutamatergic principle neurons and inhibitory GABAergic “interneurons” (Pelkey et al., 2017). It is possible that different divisions or cell types were preferentially impacted in the above studies and generated different behavioral phenotypes. To test this possibility, we focused on the GABAergic outputs from the hippocampus to examine their specific contributions to the regulation of locomotion. We found that among hippocampal GABAergic outputs, those to the septum were particularly engaged in locomotion regulation. Consistent with earlier studies showing the importance of the hippocampo-septal pathway (Bender et al., 2015), this study reveals a new hippocampal mechanism for locomotion regulation.

## Results

### Activation of septum-projecting hippocampal interneurons decreases locomotion and exploratory behavior

Recently, inhibitory cells in the dorsal hippocampus were shown to play an important role in locomotion (Arriaga & Han, 2017; Gois & Tort, 2018). The activities of somatostatin-expressing and parvalbumin-expressing inhibitory cells correlate with locomotion (Arriaga & Han, 2017; Geiller et al., 2020). Although commonly referred to as “interneurons”, many GABAergic inhibitory neurons in the hippocampus send long-range projections to extrahippocampal regions (Klausberger & Somogyi, 2008). Here, we focused on the role of these hippocampal inhibitory outputs in locomotion regulation. We first examined the distribution of the long-range hippocampal GABAergic projections (Figure. 1a, b). We injected an adeno-associated virus (AAV) vector into the dorsal hippocampus to express EGFP under Dlx promoter for selective expression in GABAergic neurons (Dimidschstein et al., 2016). The distribution of the EGFP-positive neurons in the hippocampus was consistent with the distribution of the GABAergic interneurons in the hippocampus (Pelkey et al., 2017). While EGFP-positive axons were observed in multiple brain regions, they showed a concentration in the medial septum (MS), consistent with earlier studies (Gulyas, Hajos, Katona, & Freund, 2003; Jinno et al., 2007; Muller & Remy, 2018) (Figure. 1c, d). To identify the septum-projecting hippocampal GABAergic interneurons, we injected into the hippocampus an AAV for Cre-dependent expression of EGFP under the Dlx promoter (AAV-Dlx-DIO-EGFP); and injected into the MS AAV2-retro-Cre, which could be uptaken by axonal terminals and retrogradely transported to the soma of the neurons (Tervo et al., 2016). (Figure 1— figure supplement1a).

**Figure 1.**
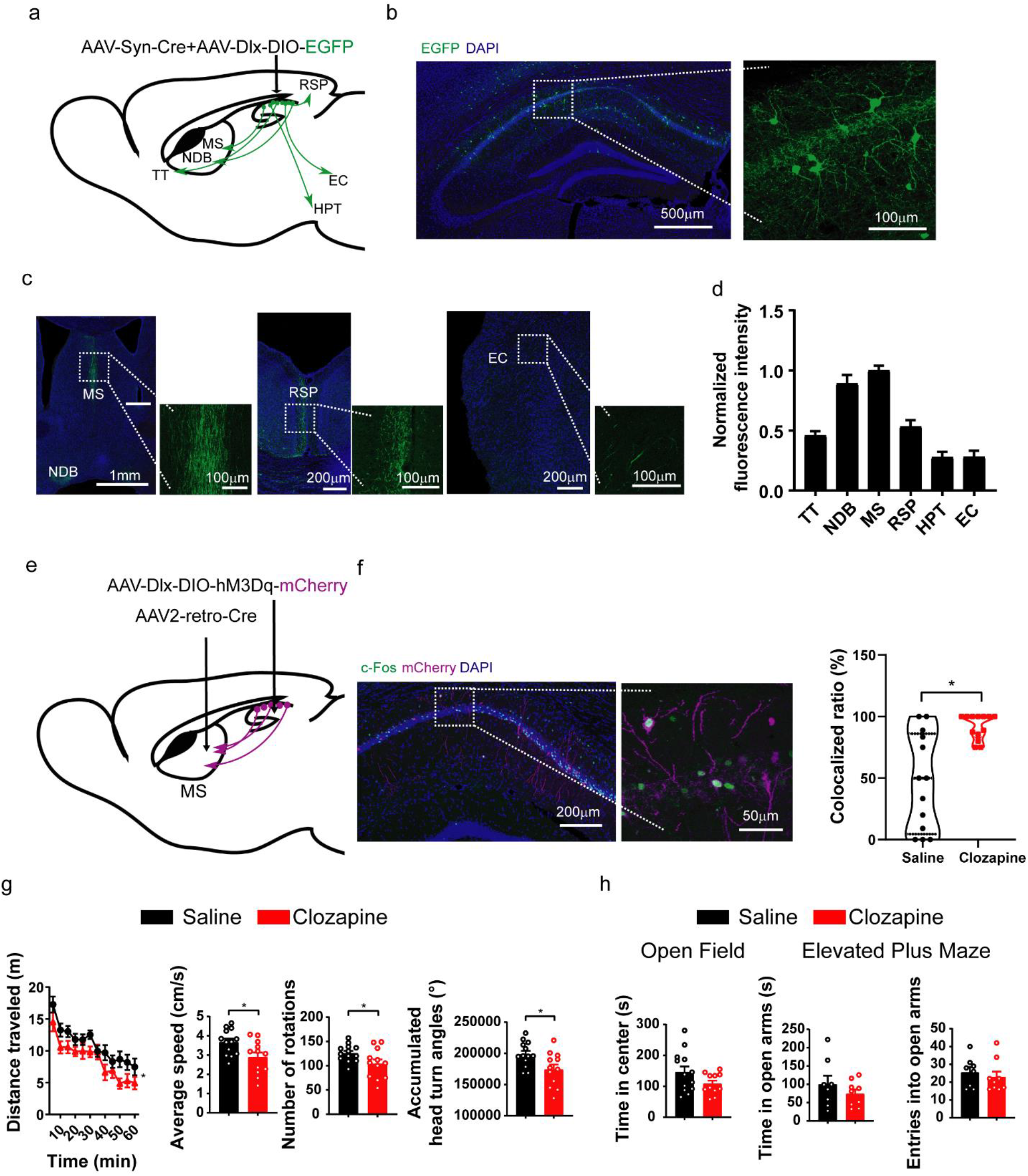
Activation of septum-projecting hippocampal GABAergic interneurons decreases locomotion and exploratory behavior. (**a**) Tracing GABAergic projections from the hippocampus with injection of AAV expressing EGFP into the hippocampus. (**b**) Expression of EGFP in hippocampal GABAergic interneurons. (**c**) Representative photos showing the distribution of EGFP-positive axonal terminals originated from the hippocampus in the medial septum nucleus (MS) and diagonal band nucleus (NDB), retrosplenial cortex (RSP), and entorhinal cortex (EC). (**d**) Quantification of the EGFP-positive axonal terminals. The hippocampal GABAergic interneurons mainly project to the MS and the NDB. (TT: n=8 sections; NDB: n=19 sections; MS: n=18 sections; RSP: n=17 sections; HPT: n=6 sections; EC: n=8 sections; from 3 mice). Data are represented as mean ± SEM. (**e**) Expression of hM3Dq in septum-projecting hippocampal GABAergic interneurons by AAV injections. (**f**) Intraperitoneal injections of clozapine (0.1mg/kg), a ligand of hM3Dq, increased neuronal activity of septum-projecting hippocampal GABAergic interneurons measured by double labeling of c-Fos (green) and mCherry (purple) (Saline, n=13 sections, total 62 cells from 2 mice; Clozapine, n=13 sections, total 73 cells from 2 mice) (Mann–Whitney U test, *p<0.05).(**g**) Activation of the septum-projecting hippocampal GABAergic interneurons with clozapine decreased mouse locomotion activities measured by the distance traveled in the open field, number of rotations and accumulated head turn angles (Two-way ANOVA, F(1,23)=6.35, p<0.05; Two-tailed t-test, *p<0.05). (**h**) Activation of the septum-projecting hippocampal GABAergic interneurons did not change time spent in the center of the open-field test (Two-tailed t-test, p=0.1), and time spent in the open arms (Two-tailed t-test, p=0.31) and the number of entries into the open arms of the elevated plus maze test (Two-tailed t-test, p=0.55) (Open field: Saline: n=13; Clozapine: n=11; Elevated plus maze: Saline: n=8; Clozapine: n=9).

The EGFP-positive neurons distributed in broad hippocampal regions but had a concentration in the str. oriens of the hippocampus region (Figure 1—figure supplement1b, c, d), consisting with previous studies (Muller & Remy, 2018).

To functionally control these septum-projecting hippocampal GABAergic interneurons, we injected AAV2-retro-Cre in the MS, and injected Cre-dependent AAV (AAV-Dlx-DIO-hM3Dq) in the hippocampus to express hM3Dq, an excitatory DREADD effector (Roth, 2016) (Figure. 1e). With immunostaining of c-Fos, an immediate early gene reporting neuronal activities, we confirmed that a low dose of clozapine [a ligand of hM3Dq (Gomez et al., 2017), 0.1 mg/kg, intraperitoneal injection] increased neuronal activity of hM3Dq-positive septum-projecting hippocampal GABAergic interneurons (Figure. 1f). We then examined the mice in behavioral tests. In open-field, the mice decreased the speed of locomotion and exploratory behavior upon the activation of the septum-projecting hippocampal GABAergic interneurons (Figure. 1g; Figure 1— figure supplement2) without a change in the time spent in the center of the open field arena, a parameter frequently used to monitor anxiety level (Figure. 1h). Similarly, the mice did not show an anxiety phenotype in elevated-plus maze test (Figure. 1h). These results indicate that septum-projecting hippocampal GABAergic interneurons negatively regulate mouse locomotion.

### Activation of hippocampal inhibitory output to the MS decreases locomotion and exploratory behavior

The septum-projecting hippocampal GABAergic interneurons may regulate locomotion through their inhibition of the local hippocampal network or through their inhibition in the MS. To determine if their axons in the MS is sufficient to regulate locomotion, we stimulated these axon terminals with locally injected clozapine-N-oxide (CNO), a selective ligand for hM3Dq, through cannula implanted into the MS (Figure. 2a). Pharmacogenetic activation of these inhibitory inputs to the MS decreased locomotion (Figure. 2b), similar to the effects produced by activating the septum-projecting hippocampal GABAergic interneurons (Figure. 1g), suggesting that through direct inhibition of axons in the MS, the septum-projecting hippocampal GABAergic interneurons negatively regulate locomotion.

**Figure 2.**
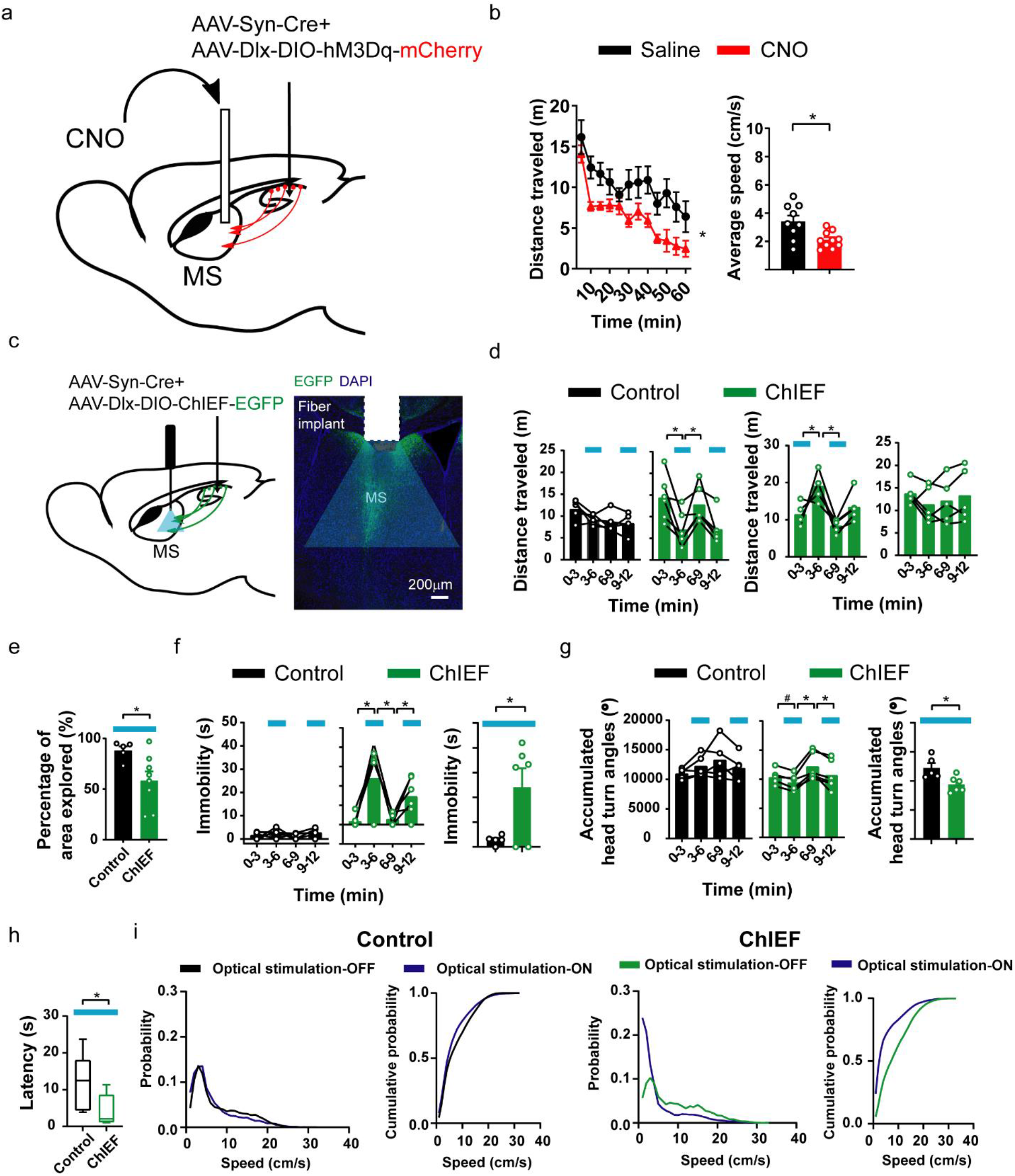
Activation of hippocampal inhibitory output to the MS decreases locomotion and exploratory behavior. (**a**) Pharmacogenetic activation of the hippocampal inhibitory inputs to the MS with local infusion of CNO in the MS. (**b**) CNO decreased locomotion in the open field measured by distance traveled (Two-way ANOVA, Group, F (1, 17) = 9.00, *p<0.05) and average speed (Two-tailed t-test, *p<0.05) (Saline, n=9 mice; CNO, n=10 mice). (**c**) Optogenetic activation of the hippocampal inhibitory inputs to the MS with blue light delivered into the MS. ChIEF, an excitatory channelrhodopsin, was expressed in the hippocampal GABAergic neurons with AAVs in the “ChIEF” group. Mice in the control group received AAVs expressed EGFP in the “Control” group. (**d-i**) Blue light delivered to the MS decreased mouse locomotion measured by distance traveled (**d**), percentage of the open field area the mice explored (**e**), immobility (**f**), accumulated angles of head turns (**g**), latency to reach a low speed (**h**), and the distribution of locomotion speed (**i**). The blue bars indicate the delivery of blue light. (**d**: Two-way ANOVA. Left: ChIEF, Light, F (1,5) = 30.42, p<0.05; Right: ChIEF, Light, F (1, 4) = 10.15, p<0.05; Tukey’s multiple comparison test, *p<0.05. **e**: Two-tailed t-test, *p<0.05. **f**: Left: Two-way ANOVA. Control, Light, F (1, 4) = 5.409, p=0.08; ChIEF, Light, F (1, 5) = 10.57, p<0.05; Tukey’s multiple comparison test, *p<0.05; Right: Two-tailed t-test, *p<0.05. **g**: Left: Two way ANOVA. Control, Light, F (1, 4) = 0.03, p=0.86; ChIEF, Light, F (1, 5) = 22.01, p<0.05; Tukey’s multiple comparison test, *p<0.05; Right: Two-tailed t-test, *p<0.05. **h**: Mann-Whitney U test, *p<0.05. **i**: Kolmogorov-Smirnov test, Control, p=0.43; ChIEF, p=0.05) (Control, n=5 mice; ChIEF, n=6 mice; (**d**) Right: ChIEF, n=5).

For temporally precise control of the projections to the MS, we expressed a channelrhodopsin, ChIEF, in hippocampal interneurons and delivered light to the MS to stimulate the GABAergic hippocampal axons distributed there (Figure. 2c). Optical stimulation acutely decreased mouse locomotor activities measured by multiple parameters including the distance traveled, the area of open field the animals explored (Figure. 2d), mobility time (Figure. 2f) and total head movement in the open field (Figure. 2g). The stimulation also decreased the latency to reach a low speed (Figure. 2h) and slightly changed the speed distribution (Figure. 2i). The decreased locomotion was reversible (Figure. 2d) and was not accompanied by a change in anxiety level (Figure 2—figure supplement1a, b, c). Consistent with above pharmacogenetic findings, these results indicate that activation of the GABAergic septal projections from the hippocampus decreases animal locomotion and exploratory behavior.

### Inhibiting inhibitory hippocampal output to the MS increases locomotion and exploratory behavior

Next, to test if the inhibitory hippocampal outputs to the MS can bidirectionally regulate locomotion, we expressed an inhibitory opsin, Jaws (Chuong et al., 2014), in the hippocampal interneurons and delivered light to the MS to silence the GABAergic axons originated from the hippocampus (Figure. 3a). This optical inhibition increased mouse locomotor activities measured by the maximum locomotion speed (Figure. 3b, e), head movement (Figure. 3c, e), the number of rearings (Figure. 3d), and habituation (Figure 3—figure supplement1a, b). Again, the increased locomotion and exploratory behavior were not accompanied by a change in anxiety level measured by the time spent in the center of the open field or time spent in the open arms of the elevated plus maze (Figure 3—figure supplement1c, d). Together, the results demonstrate that the activities of the GABAergic septum projection from the hippocampus bi-directionally regulated mouse locomotion and exploratory behavior.

**Figure 3.**
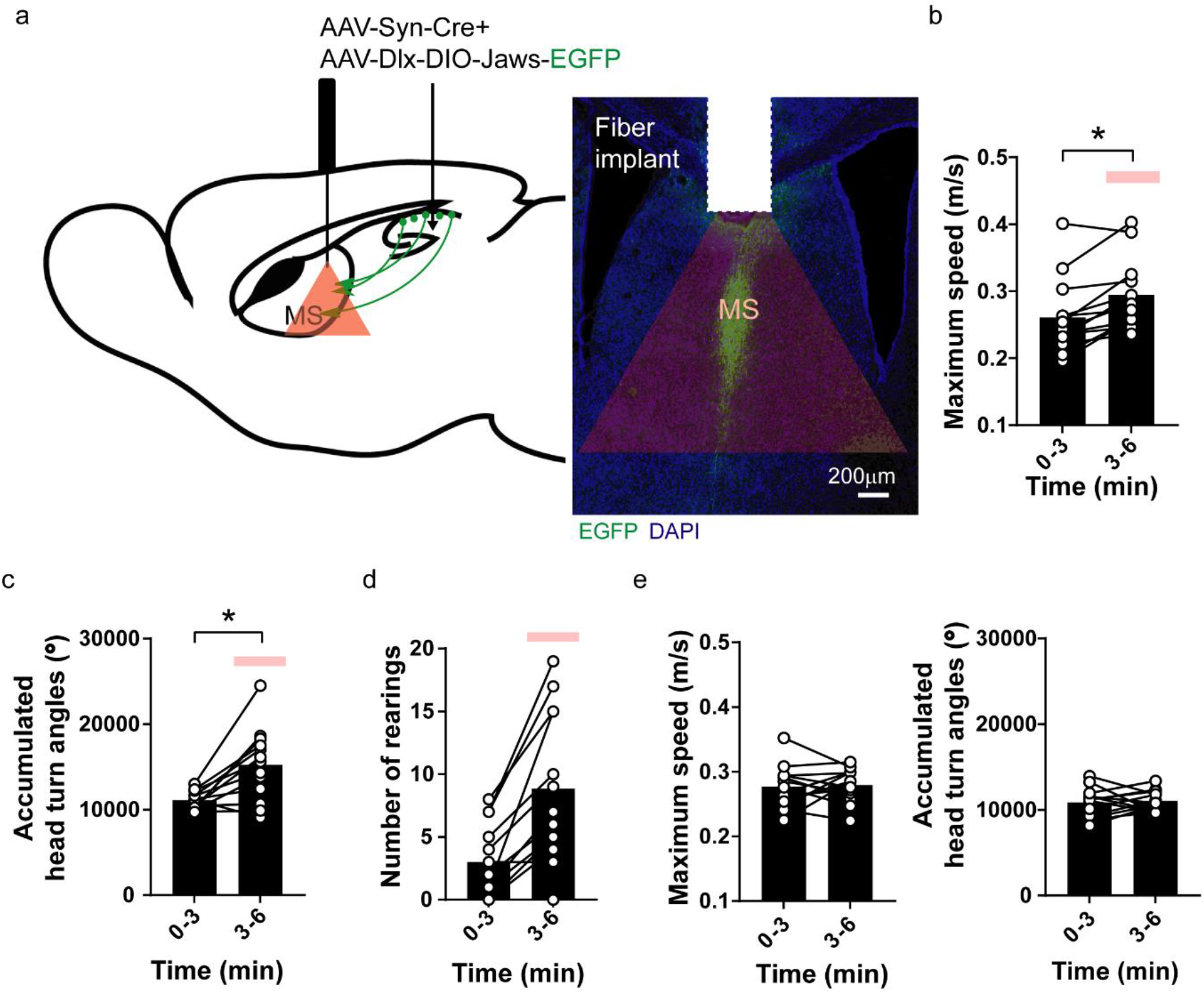
Inhibiting inhibitory hippocampal output to the MS increases locomotion and exploratory behavior. (**a**) Optogenetic inhibition of the hippocampal inhibitory inputs to the MS with blue light delivered into the MS. Jaws, an inhibitory opsin, was expressed in the hippocampal GABAergic neurons with AAVs. (**b-d**) Red light delivered to the MS increased mouse locomotion measured by maximum speed (**b**), accumulated head turn angles (**c**) and the number of rearing events (**d**) in the open field test. (Two-tailed paired t-test, *p<0.05, **p<0.001) (Dlx-Jaws, n=13 mice). The red bars indicate the delivery of optostimulation (**e**) No change in maximum speed (left) or accumulated head turn angles (right) when the optical stimulation was not delivered (Left: Two-tailed paired t-test, p=0.77; Right: Two-tailed paired t-test, p=0.71) (Dlx-Jaws, n=13 mice).

### Activation of cortex-projecting hippocampal interneurons did not change locomotor activity

In addition to the MS, the retrosplenial cortex (RSP) receives significant amount of GABAergic inputs from the hippocampus (Klausberger & Somogyi, 2008; Witter, 2010) (Figure 1c, d). Recently, we found that a group of GABAergic interneurons located at stratum lacunosum-moleculare (SLM) of the hippocampus, which express a marker gene neuron-derived neurotrophic factor (NDNF), sent long-ranged projection exclusively to the RSP (Figure. 4a, b). To determine if the RSP-projecting hippocampal interneurons also regulate locomotion, we expressed hM3Dq selectively in these NDNF-expressing interneurons by injecting Cre-dependent AAV into the hippocampus of NDNF-Cre mouse line. Despite that pharmacogenetic stimulation of these NDNF-cells significantly changed learning and memory behaviors (Guo et al., 2021), it did not alter mouse locomotor (Figure. 4c). These results demonstrated the functional heterogeneity of the hippocampal interneurons in regulation of locomotion (Figure 4—figure supplement1).

**Figure 4.**
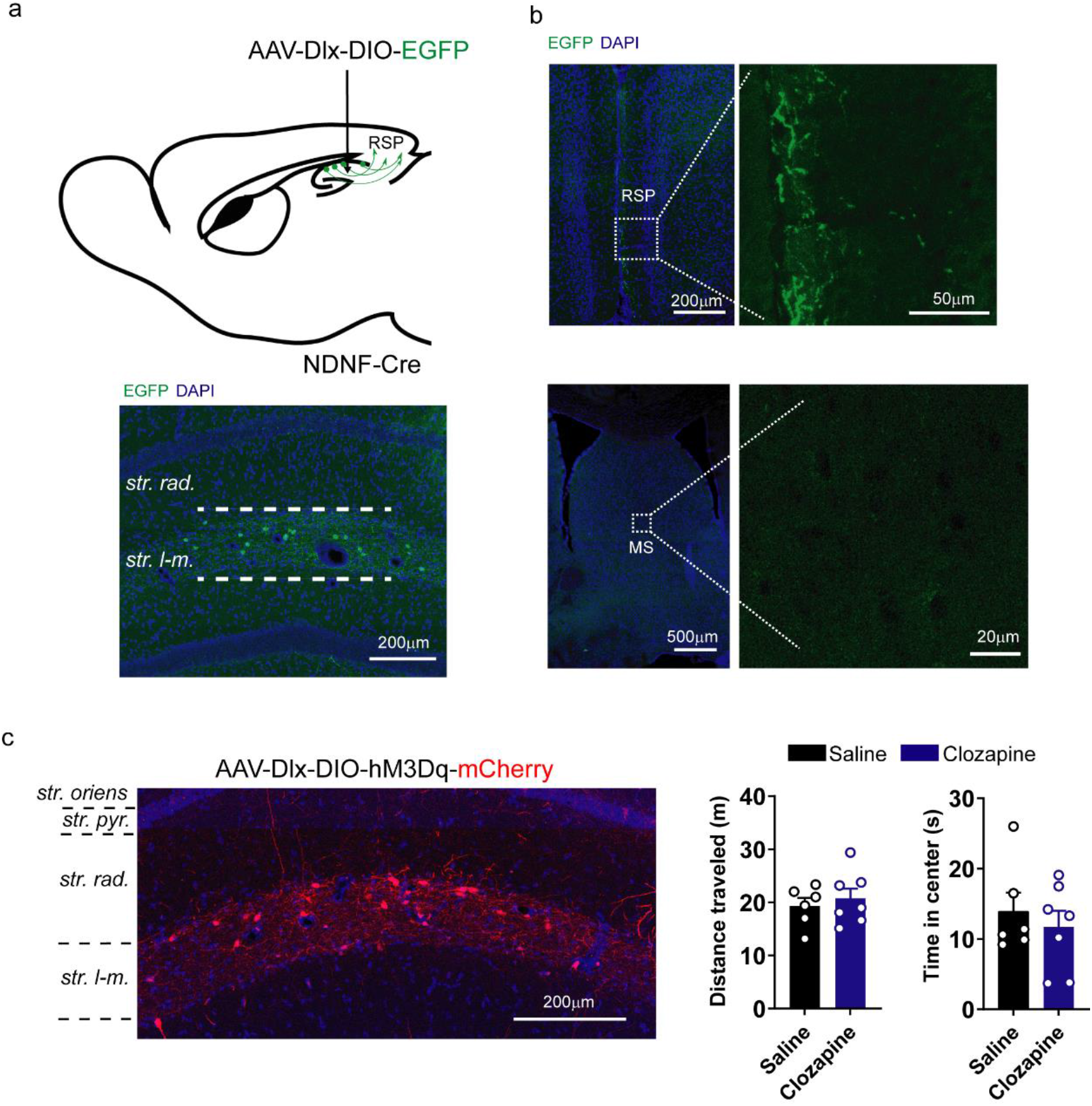
Activation of cortex-projecting hippocampal interneurons does not change locomotor activity. (**a**) Expression of EGFP in NDNF-positive interneurons in NDNF-Cre mice by injecting AAV-Dlx-DIO-EGFP into the hippocampus. (**b**) NDNF-positive interneurons in the hippocampus projected to the RSP but not to the MS. (**c)** Expression of hM3Dq in NDNF-positive interneurons in the hippocampus. (**d**) Pharmacogenetic activation of NDNF-positive interneurons with clozapine did not alter distance traveled or time spent in the center of the open field test (distance traveled: Two-tailed t-test, p=0.58; time in the center: Two-tailed t-test, p=0.53; Data are represented as mean ± SEM).

## Discussion

In brain systems, the hippocampus sits at the top of the hierarchy for sensory information processing (Squire, Stark, & Clark, 2004). The associative cortices feed multi-modal sensory information to the hippocampus to construct an integrated representation and record of the world that include the “where, when and what” information of our experiences (Eichenbaum, 2004). The information as to animal’s own locations and actions is essential for the construction and update of this neuronal representation (Gois & Tort, 2018). It is therefore not surprising that hippocampal neuronal activities closely correlate with locomotion. However, it is not clear if and how the hippocampus directly regulates locomotion through interacting with brain motor systems, and what the functional importance of this regulation is.

Addressing this question will help us to understand not only the neuronal control of locomotion but also the pathogenesis and treatment of the neuropsychiatric disorders showing functional abnormalities in the hippocampal circuits. One prominent example is schizophrenia. Anatomical, biochemical and functional alterations in the hippocampus are the most consistent findings in the patients of schizophrenia (Tamminga, Stan, & Wagner, 2010). Besides cognitive symptoms, the patients frequently showed motor symptoms, exhibiting as either dyskinesia or parkinsonism (Walther & Strik, 2012) and deficits in sensory-motor gating (Braff & Geyer, 1990; Mena et al., 2016). In animal models of schizophrenia, including both the pharmacological and genetic models, hyperlocomotion is the most common behavioral phenotype and commonly considered as a correlate of the positive symptoms (van den Buuse, 2010). Reduction of hyperlocomotion is frequently used as a behavioral readout for screening antipsychotic medicine (Powell & Miyakawa, 2006; Peleg-Raibstein, Knuesel, & Feldon, 2008; van den Buuse, 2010; Wolff et al., 2018). Determining the hippocampal mechanisms in locomotion regulation will elucidate the origin of the locomotor symptoms in patients and help us to determine the predict value of the locomotion parameters in animal models for therapeutic treatments of patients.

In the previous studies, multiple methods or protocols were used to examine the role of the hippocampus animal locomotion. The methods include activity chamber (Pierce & Kalivas, 2007; Thomas, Marcaletti, & Feige, 2011), open field (Seibenhener & Wooten, 2015), treadmill (Leblond, L’Esperance, Orsal, & Rossignol, 2003), and running wheel (Thomas et al., 2011). The duration of recordings of animal activity ranged from five minutes to several hours (Pierce & Kalivas, 2007; Thomas et al., 2011). This variety of experimental conditions led to inconsistent conclusions (Douglas & Isaacson, 1964; McNaughton et al., 1983; Swerdlow et al., 2001; Degoulet et al., 2008; Kropff, Carmichael, Moser, & Moser, 2015). Although the glutamatergic projections from the MS to hippocampal interneurons, or the hippocampal excitatory outputs to the lateral septum, were shown to control locomotion (Bender et al., 2015; Fuhrmann et al., 2015), there are contradictory results as to the roles of these pathways in locomotion (McGlinchey & Aston-Jones, 2018).

In this study, we identified the inhibitory hipocampo-septal projection as a key pathway for locomotion regulation. The pharmacogenetic and optogenetic techniques allowed us to isolate this pathway for precise and reversible functional manipulations. The results indicated the GABAergic hippocampo-septal projection bidirectional and reversible regulates locomotor activities. This reveals a new function of the extrahippocampal projection of the hippocampal interneurons and builds a foundation for further elucidation of hippocampal regulation of motor activities. The MS is innervated by and projects to multiple brain regions. It is engaged in learning, memory, emotional reaction, defensive behaviors and sensorimotor gating (Tsanov, 2017, 2018; Jin et al., 2019). However, anatomically and functionally it is most closely coupled with the hippocampus (Muller & Remy, 2018; Iyer & Tole, 2020). The hippocampo-septal loop is particularly critical for the generation of hippocampal oscillations that arises during locomotion and active exploration in the environment (O’keefe & Nadel, 1978; Buzsaki, 2002; Drieu & Zugaro, 2019). MS plays a pacemaker role for hippocampal theta oscillation (Buzsaki, 2002; Tsanov, 2017). Our data have not touched the question that if these GABAergic projections to the septum exerted their locomotion regulation effects through altering functions of the hippocampo-septal loop, or act on the motor structures downstream of the septum. Besides the hippocampus, the MS projects to the cortex, thalamus, and midbrain structures which may serve as an interface between the hippocampus and the motor control systems. Recently, we developed techniques for stepwise polysynaptic tracing and genetic control (Du et al, manuscript under submission; Li et al., manuscript in press). These techniques allow us to selectively control the septal neurons innervated by the hippocampal GABAergic outputs and their postsynaptic neurons distributed in the brain regions outside the hippocampo-septal systems. With these techniques we may be able to identify the next stage of the hippocampo-septal pathway in regulating locomotion.

## Material and Methods

### Animal

8 to12-week-old C57BL/6J (B6J) male mice were obtained from UT Southwestern animal breeding core or The Jackson Laboratory. We used heterozygotes (+/−) NDNF-Cre transgenic male mice and maintained on a C57BL/6J background (The Jackson Laboratory). Animal work was approved and conducted under the oversight of the UT Southwestern Institutional Animal Care and Use Committee and complied with Guide for the Care and Use of Laboratory Animals by National Research Council.

### Stereotaxic surgery and adeno-associated virus (AAV) injection

Mice were anesthetized with intraperitoneal (i.p.) injection of Tribromoethanol (Avertin) (125-250 mg/kg) before the stereotaxic surgery or anesthetized with 1-3% isoflurane and placed in a stereotaxic instrument (Kopf Instruments). Viral vector construction and preparation were the same as previously described (Guo et al., 2021). To identify the septum-projecting hippocampal GABAergic interneurons, AAV2-retro-Cre (0.75 μl) was injected into the septum (coordinates: A/P 0.80 mm, M/L ±0.00 mm, D/V 4.20 mm) and infused slowly over 7.5 minutes (rate: 0.1μl/minute) using a microdriver with a 10μl Halmiton syringe connected to a glass pipette, then AAV-Dlx-DIO-EGFP was injected into the CA1 (coordinates A/P −1.95 mm, M/L ±1.25 mm, D/V 1.25 mm). To manipulate the septum-projecting hippocampal GABAergic interneurons, AAV2-retro-Cre (0.75 μl) was injected into the septum (coordinates: A/P 0.80 mm, M/L ±0.00 mm, D/V 4.20 mm) and infused slowly over 7.5 minutes (rate: 0.1μl/minute), then AAV-dlx-DIO-hM3Dq-mCherry was injected into the CA1 (coordinates A/P −1.95 mm, M/L ±1.25 mm, D/V 1.25 mm).

To optogentically activate the hippocampal inhibitory output to the septum, the virus AAV-Dlx-DIO-ChIEF-EGFP and AAV-Syn-Cre were mixed in the ratio 4:1 (To optically inhibit the hippocampal inhibitory output to the septum, we used the virus mixture of AAV-Dlx-DIO-Jaws-EGFP and AAV-Syn-Cre). Viruses (0.5 μl for each target) infused slowly over 5 minutes (rate: 0.1μl/minute) into the CA1 (coordinates A/P −1.95 mm, M/L ±1.25 mm, D/V 1.25 mm) bilaterally using a microdriver with a 10μl Halmiton syringe connected to a glass pipette, after the virus injection, flat-cut 400 μm diameter optic fiber with ferrule (Ø 400 μm; CFM14U-20, Thor labs) was implanted on the top of the medial septum (A/P 0.60 mm, M/L 0.00 mm, D/V 2.50 mm) and cemented in place using dental cement and C&B-Metabond (Patterson dental, MN).

To pharmacogenetically manipulate the hippocampal output to the septum, the virus AAV-dlx-DIO-hM3Dq-mCherry and AAV-Syn-Cre were mixed in the ratio 4:1 (0.5 μl for each target) using the same stereotaxic coordinates to target the CA1. After the virus injection, a gauge 28 guide-cannula was implanted on the top of the medial septum (A/P 0.70 mm, M/L 0.00 mm, D/V 2.50 mm) and cemented in place using dental cement and C&B-Metabond (Patterson dental, MN).

To pharmacogenetically manipulate the NDNF-expressing interneurons in the hippocampus, we bilaterally injected 0.5 μl of AAV hDlx-DIO-hM3Dq-mCherry in the dorsal hippocampus (A/P: −1.95, mm, M/L: +1.25 mm, D/V: 1.45 mm) of NDNF-Cre+ and Cre littermates.

### Optogenetic and pharmacogenetic manipulation

For optogenetics in ChIEF experiment, blue laser was delivered through a fiber optic cord using a DPSS Blue 473 nm Laser source (MBL-III-473/ 1~100mW, Opto Engine LLC). A train of blue laser pulses (10mW, 20Hz, 10ms duration, 40ms interval) was generated and controlled by Optogenetics Pulser (Prizmatix). In Jaws experiments, continuous orange-red LED was delivered through a fiber optic cord using a high power Orange-Red ~625nm LED module at 10mW (Prizmatix). Light intensity was calibrated with the PM100D Console (Thorlabs). For pharmacogenetic experiments, i.p. injections of clozapine (0444, Tocris Bioscience) (0.1mg/kg) or control vehicle saline (0.9% NaCl) were administered 30 minutes before the behavioral test. For terminal manipulation experiments, intracranial injections of 300nL clozapine (0.001mg/mL) or control vehicle saline (0.9% NaCl) were administered 15 minutes before the behavioral test.

### Open field test

Animals were handled for 1-2 minutes a day for seven days prior to the open field test. The open field apparatus was a custom-made testing 50 × 50 cm chamber. A video camera was placed above the open field and mice trace were tracked using ANY-maze video tracking system. Mice were place in the center of the open field area prior to initiation of tracking. The center of the open field was defined as 20 × 20 cm square in the geometric center of the arena. Each chamber was cleaned between individual animal testing. To calculate the percentage of the open field area each mouse explored, the open field arena was divided into 100 grids (5 × 5 cm). If the mouse passed through the corresponding grid, then the grid would be counted as being explored. The exploration percentage of the open field ranges from 1 to 100%.

### Elevated plus maze (EP)

Animals used in the elevated plus maze (EPM) were tested in the open field before. The EPM apparatus was elevated 38.7 cm above the floor and consisted of two open arms (30.5 cm in length; 5 cm in width) and two closed arms (30.5 cm in length; 6 cm high wall; 5 cm in width). Open arms and closed arms are all connected to a center platform in the middle (5 cm in length and width). The behavior was recorded and analyzed by ANY-maze video tracking system.

### Tissue processing, immunohistochemistry, and cell counting

For regular preparation with no need to do immunohistochemical staining, mice were anesthetized by an intraperitoneal (i.p.) injection of Tribromoethanol (Avertin) or anesthetized with 1-3% isoflurane and were perfused with phosphate buffered saline (PBS) followed by 4% paraformaldehyde (PFA) in PBS. Brains were post-fixed in 4% PFA overnight and were cryoprotected in 30% sucrose. Brains were cut into 40 μm sections on a cryostat (Leica CM1950) and were collected in PBS and stored at 4 degrees Celsius. Finally, sections were then mounted on slides and stained with DAPI. Sections were imaged on Zeiss LSM 880 confocal microscope with a 5x, 10x, and 20x objective under control of Zen software.

For c-Fos immunohistochemistry staining, mice were injected with clozapine (0.1mg/kg) and then transferred to a new clean cage and single housed for 1 hr before the perfusion. Brains were post-fixed in 4% PFA overnight and were cryoprotected in 30% sucrose. Brains were cut into 30 μm sections on a cryostat (Leica CM1950) and were collected in PBS. Sections were washed in PBS and blocked in 10% horse serum, 0.2% bovine serum albumin (BSA) and 0.5% Triton X-100 in PBS for 2 hrs at room temperature. For immunohistochemistry staining, sections were incubated overnight in primary antibodies (anti-cFos antibody: 1:1000, catalog # 226 003, Synaptic Systems (SYSY)) with 1% horse serum, 0.2% BSA and 0.5% Triton X-100 in PBS at 4°C. Sections were washed in PBS and reacted with fluorescent secondary antibodies (goat anti-rabbit Alexa Fluor 488, 1:500, Invitrogen, catalog # A-11034) in 1% horse serum, 0.2% BSA and 0.5% Triton X-100 in PBS for 2 hrs at room temperature. Sections were then mounted on slides and stained with DAPI. Sections were imaged on Zeiss LSM 880 confocal microscope with a 5x, 10x, and 20x objective under control of Zen software.

To quantify the number of activated hM3Dq expressing septum-projecting inhibitory cells in the hippocampus, the hippocampus were outlined as regions of interest (ROIs) and the colocalized ratio was calculated as ((cFos^+^ & mCherry^+^) / (mCherry^+^)) × 100. To quantify the fluorescence intensity of the hippocampal projections in each innervated regions, mean intensity of each ROI was acquired using ZEISS ZEN Microscope Software. And the normalized fluorescence intensity was calculated as ((mean fluorescence intensity) / (average mean fluorescence intensity of the medial septum)).

### Statistical analysis

Data are presented as means ± SEM and all statistical analyses of the data were performed using GraphPad Prism software (GraphPad Software Inc., La Jolla, USA). Student’s unpaired t-test was used to analyze two independent samples and paired t-test was used to analyze two dependent samples. To test the optical stimulation effects in two groups, open field results were analyzed by two-way repeated measures ANOVA with “Order” and “Light” as within-subject factors followed by multiple comparisons tests. 1hr (5-minutes bin) open field was analyzed by two-way repeated measures ANOVA with time (minutes) as a within-subject factors and “Group” as a between-subject followed by multiple comparisons tests. Kolmogorov-Smirnov test was used to compare the speed distribution of two groups. Mann–Whitney U test was used to compare c-Fos activity (Figure 1f) and speed latency (Figure 2h). A p-value less than 0.05 (*p < 0.05, **p<0.001) was considered statistically significant.

## Acknowledgements

We thank Elizabeth Li and Wenqin Rita Du for help with AAV preparation and production. We thank Dr. Brad Pfeiffer, Dr. Julian Meeks, and Dr. Carol Tamminga for useful comments.

