## Supplementary figures for "Inhibitory hippocampo-septal projection controls locomotion and exploratory behavior"

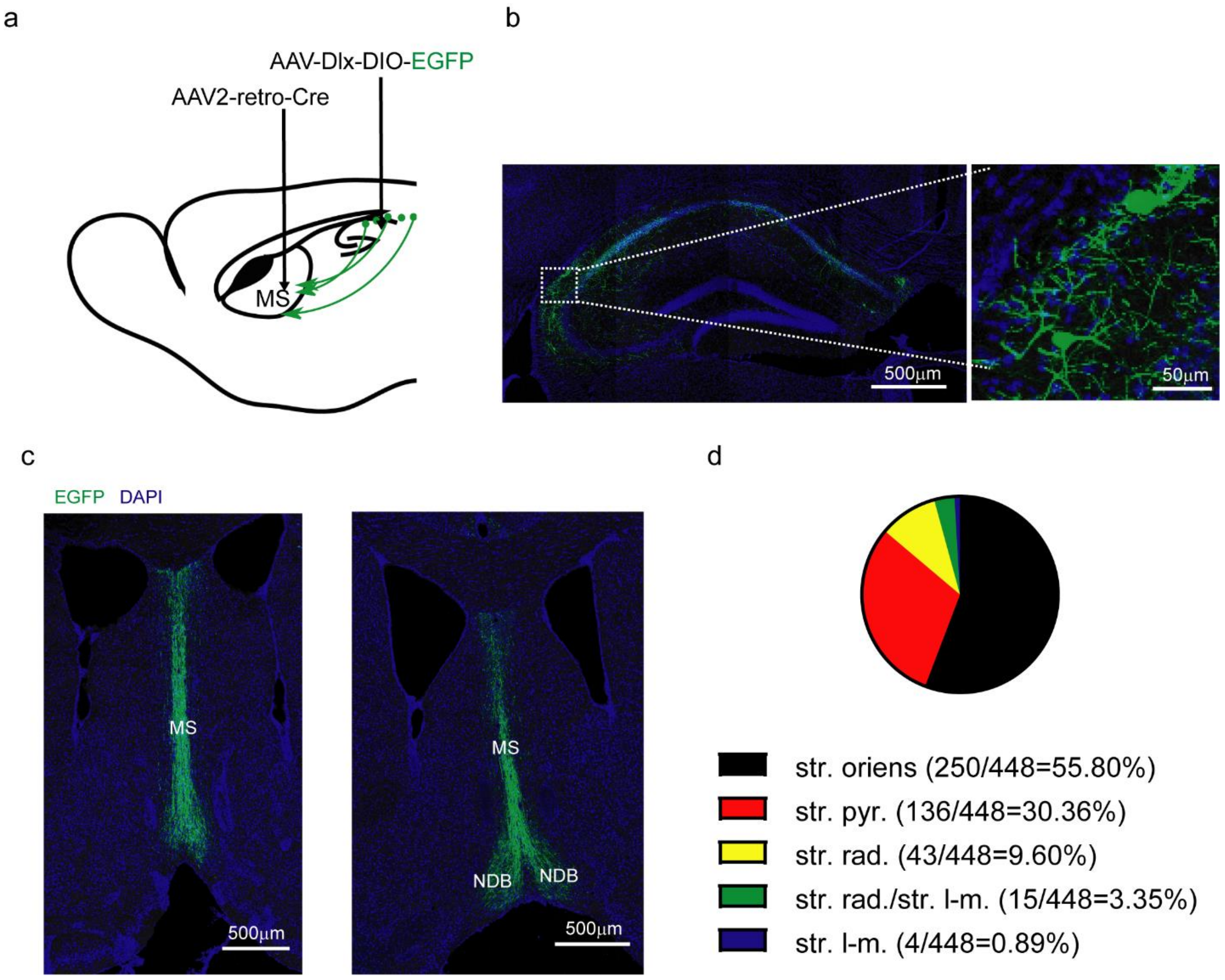


**Figure 1—figure supplement 1.** Characterizing the septum-projecting hippocampal GABAergic interneurons. (**a**) To label the septum-projecting hippocampal GABAergic interneurons, a retrograde-Cre virus (AAV2-retro-Cre) was injected into the septum and an AAV expressing a Cre-dependent EGFP reporter under the control of the Dlx enhancer was injected into the hippocampus. (**b**) EGFP-labeled septum-projecting hippocampal GABAergic interneurons were labeled in the str. oriens, str. pyr., str. rad, and str. l-m of the hippocampus. (**c**) EGFP-labeled hippocampal GABAergic interneuron terminals were observed in the MS and NDB. (**d**) The distribution ratio of septum-projecting hippocampal GABAergic interneurons (Total labeling 448 cells in 95 slices from 3 mice) in different layer of the hippocampus. Most of the septum-projecting hippocampal GABAergic interneurons were located in the str. oriens (250 cells) of the hippocampus.

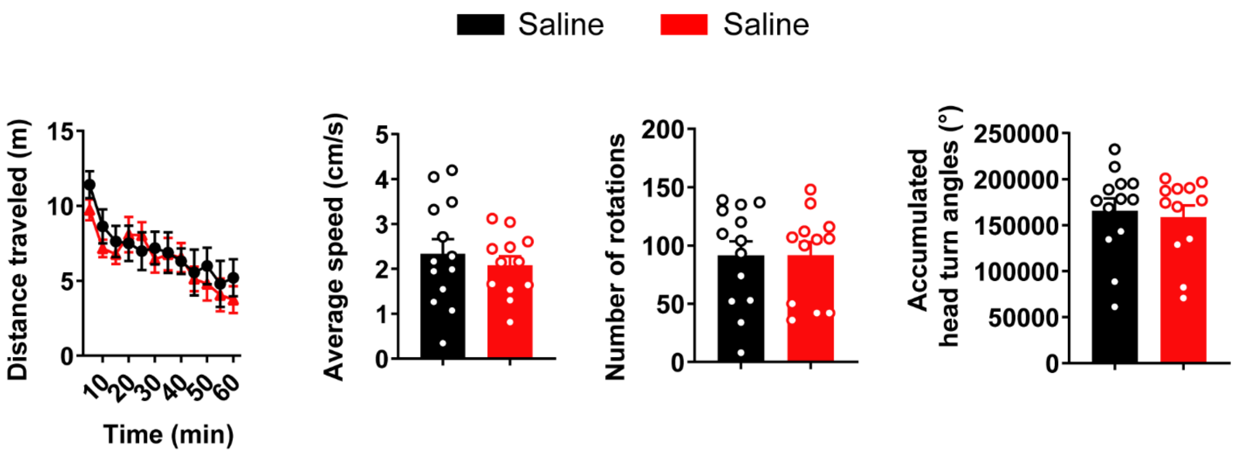


**Figure 1—figure supplement 2.** Two groups do not show difference when the septum-projecting hippocampal GABAergic interneurons were not activated (distance traveled: Two-way ANOVA, F(1,23)=0.24, p=0.63; number of rotations: Two-tailed t-test, p=0.99; accumulated head turn angles: Two-tailed t-test, p=0.71).

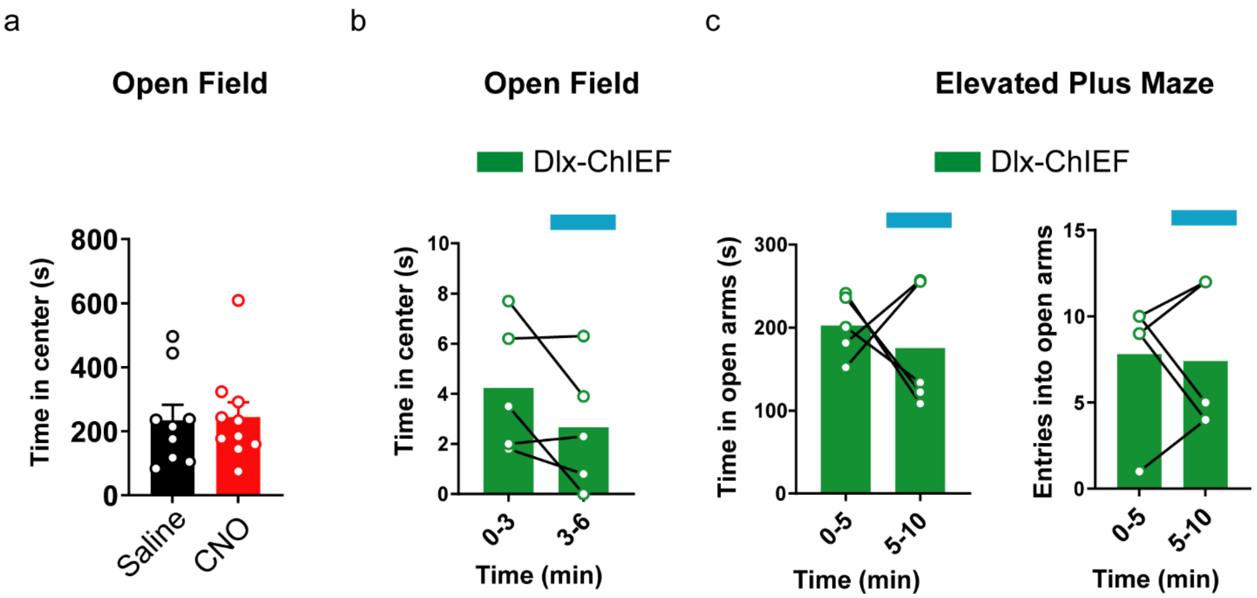


**Figure 2—figure supplement 1.** Activation of the hipppocampal inhibitory inputs to the MS does not change anxiety. (**a**) Pharmacogenetic activation of the hippocampal inhibitory inputs to the MS did not change animals time spent in the center of the open field (Two-tailed t-test, p=0.89) (GroupA, n=9 mice; GroupB, n=10 mice). (**b**) Optogenetic activation of the hippocampal inhibitory inputs to the MS did not change time spent in the center of the open field (Two-tailed paired t-test, p=0.15) (n=5 mice). (**c**) Optogentic activation of the hippocampal inhibitory inputs to the MS did not change time spent in the open arms (Two-tailed paired t-test, p=0.61,) or the number of entries into the open arms of the elevated plus maze (Two-tailed paired t-test, p=0.84) (n=5 mice).

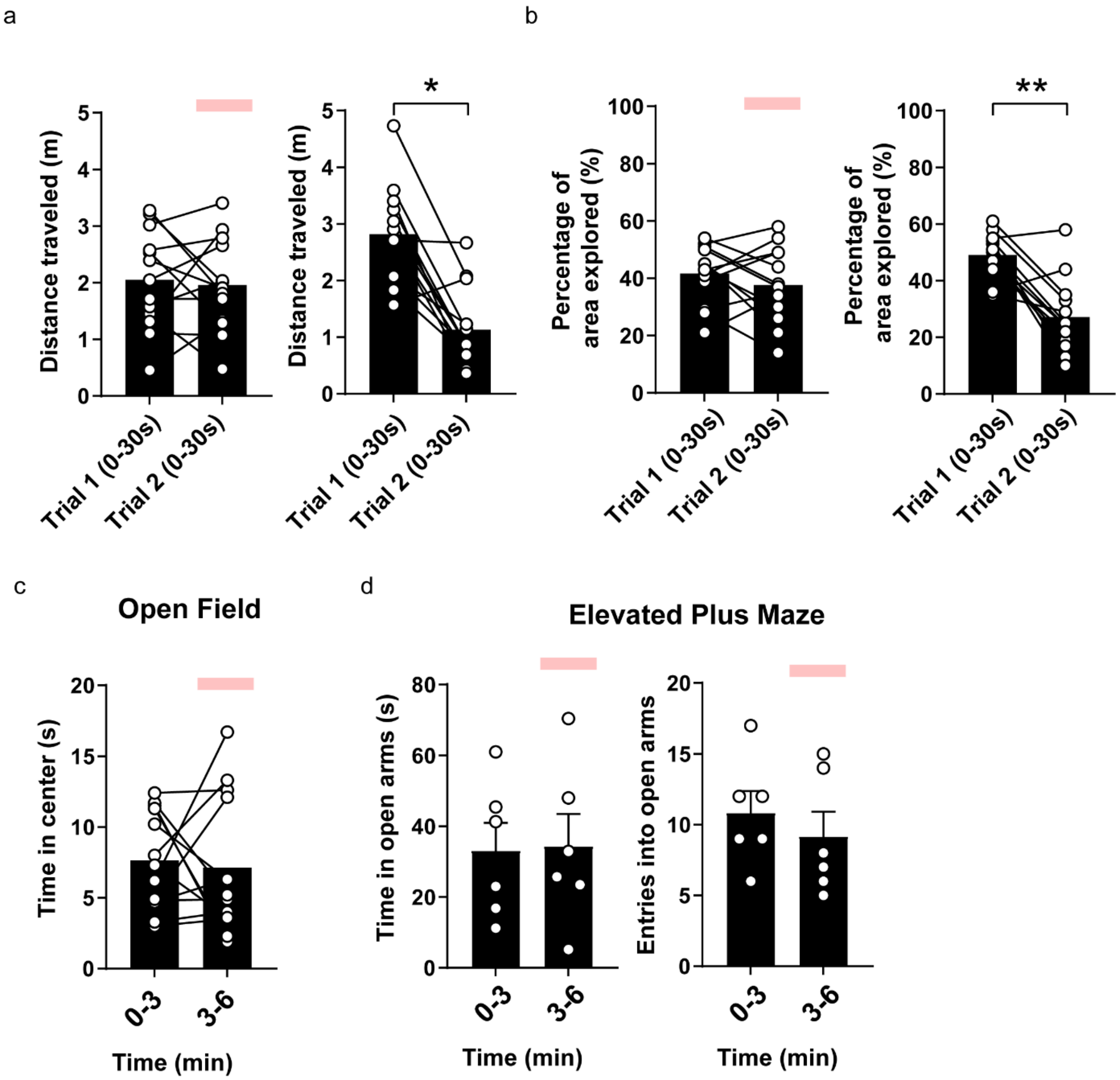


**Figure 3—figure supplement 1.** Optogenetic inhibition of the hippocampal inhibitory inputs to the MS disrupts locomotion habituation but does not change anxiety. (**a**) Optogenetic inhibition of the hippocampal inhibitory inputs to the MS disrupted locomotion habituation (Two-tailed paired t-test, p=0.72, *p<0.05). (**b**) Optogenetic inhibition of the hippocampal inhibitory inputs to the MS maintained the percentage of the open field area animals explored (Two-tailed paired t-test, p=0.29, *p<0.05). The area of the open field the animals explored decreased when the optical stimulation was not delivered (**c**) Optogenetic inhibition of the hippocampal inhibitory inputs to the MS did not change the animals time spent in the center of the open field (Two-tailed paired t-test, p=0.77). (**d**) Optogenetic inhibition of the hippocampal inhibitory inputs to the MS did not change the animals time spent in the open arms (Two-tailed paired t-test, p=0.87) and number of entries into the open arms of the elevated plus maze (Two-tailed paired t-test, p=0.48) (Dlx-Jaws, n=13 mice).

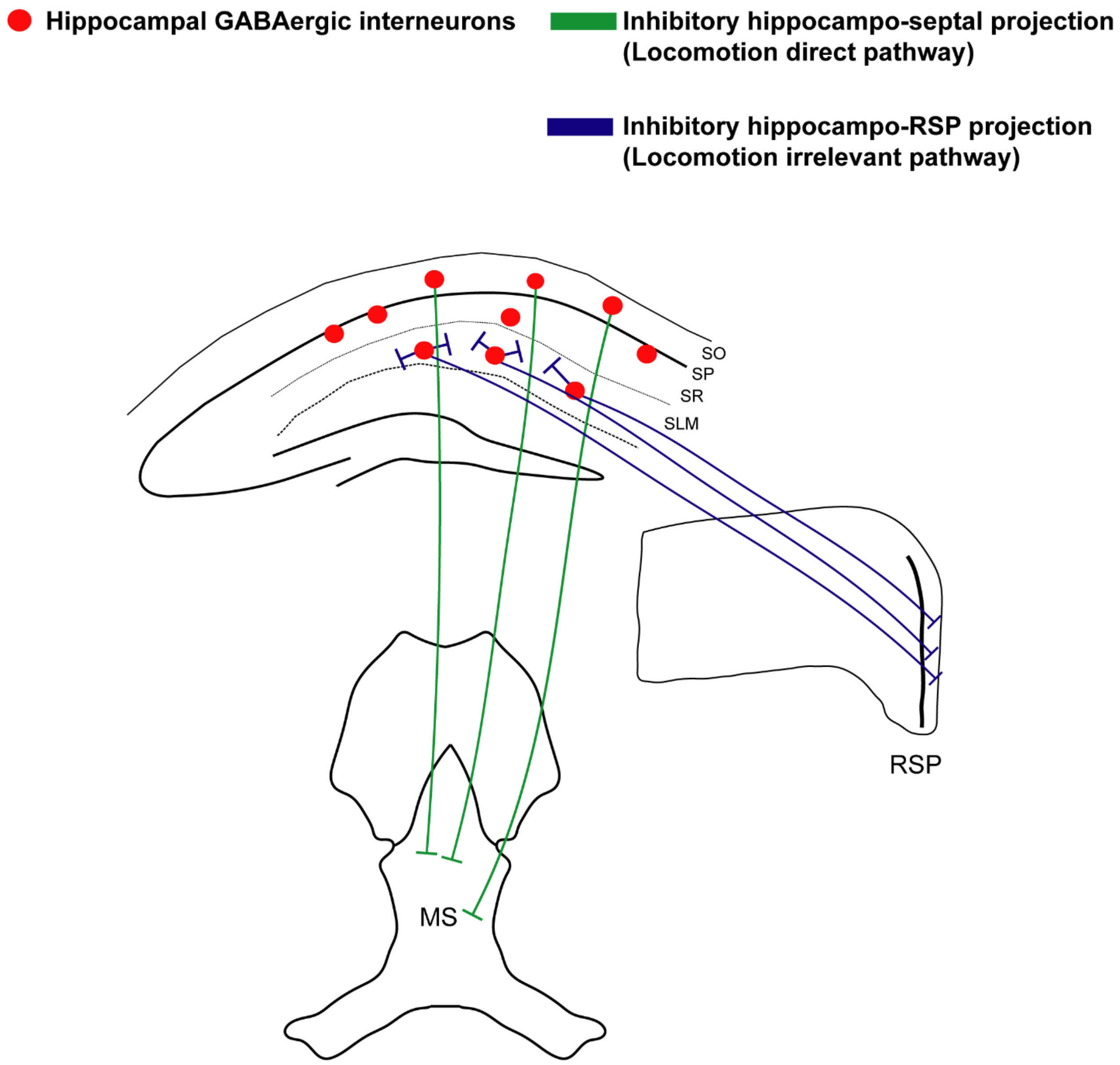


**Figure 4—figure supplement 1.** Schematic of locomotion regulation of hippocampal GABAergic interneurons. The inhibitory hippocampo-septal projection directly regulates locomotion. On the other hand, another hippocampal output pathway like hippocampo-retrosplenial cortex inhibitory pathway does not regulate locomotion.
